# cognac: rapid generation of concatenated gene alignments for phylogenetic inference from large whole genome sequencing datasets

**DOI:** 10.1101/2020.10.15.340901

**Authors:** Ryan D. Crawford, Evan S. Snitkin

## Abstract

The quantity of genomic data is expanding at an increasing rate. Tools for phylogenetic analysis which scale to the quantity of available data are required. We present cognac, a user-friendly software package to rapidly generate concatenated gene alignments for phylogenetic analysis. We applied this tool to generate core gene alignments for very large genomic datasets, including a dataset of over 11,000 genomes from the genus *Escherichia* containing 1,353 genes, which was constructed in less than 17 hours. We have released cognac as an R package (https://github.com/rdcrawford/cognac) with customizable parameters for adaptation to diverse applications.

## Background

Phylogenetic analysis is becoming an increasingly integral aspect of biological research with applications in population genetics, molecular biology, structural biology, and epidemiology ^1^. Generating a quality multiple sequence alignment (MSA) is fundamental to robust phylogenetic inference. MSA is a foundational tool in many disciplines of biology, which aims to capture the relationships between residues of related biological sequences, and therefore facilitate insights into the evolutionary or structural relationships between the sequences in the alignment.

The first analysis incorporating genetic sequences to understand the evolutionary history of an organism was a sample of 11 *Drosophila melanogaster* Adh alleles in 1983 ^2^. Since then, there has been a growing interest in using gene sequences to estimate the evolutionary relationships between organisms. However, it was quickly observed that individual gene trees are often inaccurate estimations of the species tree ^3^. These incongruencies can arise from errors while building the tree, or from biological processes such as incomplete lineage sorting, hidden parology, and horizontal gene transfer ^4^.

One approach for mitigating the incongruence between gene and species trees is the analysis of multiple genes at multiple loci concatenated into a supergene to generate more precise phylogenies ^5–9^. This approach better leverages the large quantity of available data using multiple genes to substantially increase the number of variant sites and minimize the stochastic errors that may be associated with the limited information contained in a single gene ^10^. This approach to infer the species tree has also been shown to be accurate under a range of simulated conditions, in spite of the biological processes which might pose a challenge to accurate phylogenetic inference ^11,12^.

Prior selection of a gene or set of genes for a given species is a commonly used strategy for selecting phylogenetic marker genes. For bacteria, the most commonly used marker gene for phylogenetic analysis is the 16S rRNA gene ^13^. This gene is ubiquitous in bacteria and archaea with highly conserved and variable regions which makes it a useful marker for estimating the evolutionary relationships between prokaryotes; however, this gene evolves slowly, often resulting in few variant positions within a species. Curated methods for selecting marker genes, such as multi-locus sequence typing, expand the number of marker genes for a given species, and have led to improved resolution within a species ^14^. However, this approach remains limited in that only a small number of curated genes are selected for a specific species, limiting its application to understudied organisms. Recently this concept has been expanded to include 400 marker genes commonly present in bacteria and archaea concatenated into a supergene for phylogenetic analysis of prokaryotes ^10^. While these tools have many useful applications, relying on a limited number of predefined genes may limit the number of phylogenetically informative markers contained in a given dataset, which is important in situations where maximizing variation to distinguish closely related isolates is required.

In this work, we present cognac (<u>co</u>re <u>g</u>e<u>n</u>e <u>a</u>lignment <u>c</u>oncatenation), a novel data-driven method and rapid algorithm for identifying phylogenetic marker genes from whole genome sequences and generating concatenated gene alignments, which scales to extremely large datasets of greater than 11,000 bacterial genomes. Our approach is robust when handling data sets with extremely diverse genomes and is capable of creating an alignment with large numbers of variants for phylogenetic inference.

## Results

### Implementation

The inputs to cognac are fasta files and genome annotations in gff format (Fig. 1). First, the sequences corresponding to the coding genes are extracted using the coordinates provided in the gff file, and the nucleotide sequence for each gene is translated. To identify phylogenetic marker genes, CD-HIT is then used to cluster the amino acid sequences into clusters of orthologous genes (COGs) by their sequence similarity and length ^17^. By default, COGs are defined at a minimum of 70% amino acid identity, and that the alignment coverage for the longer sequence is 80% at minimum.

**Fig. 1.**
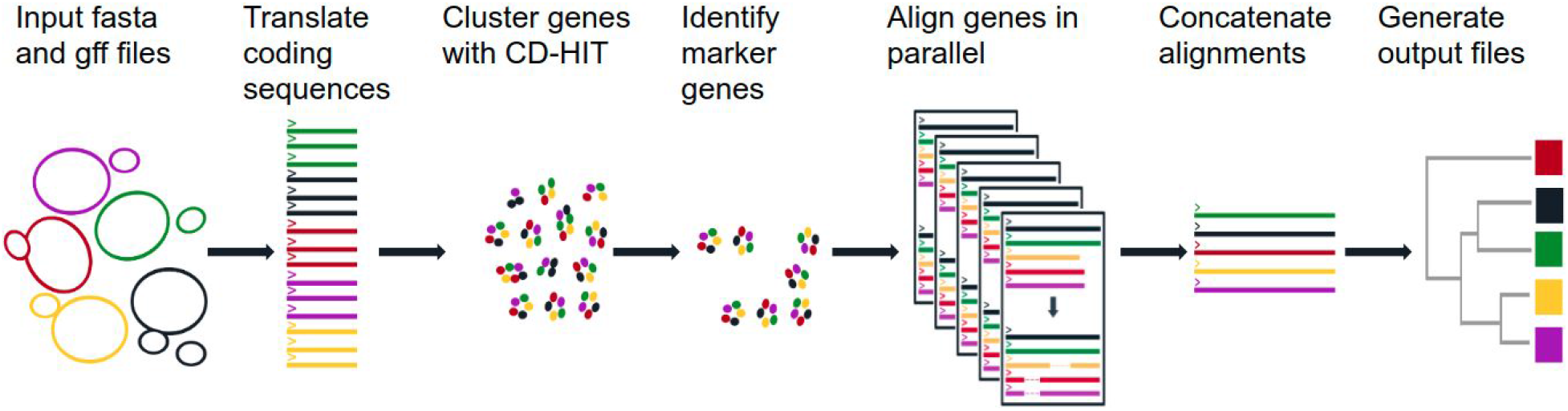
Overview of the cognac algorithm. Whole genome sequences and gene annotations are input, and the coding sequences are extracted and translated to return the amino acid sequences. The amino acid sequences are clustered to identify orthologous genes and the single copy, core genes are extracted from the dataset. For each core gene, unique alleles are identified and aligned and the alignment is parsed to represent the aligned sequence for the full dataset. Alignments are then concatenated, and are ready for downstream analysis.

The CD-HIT output file is then parsed and marker genes within the dataset are selected for inclusion in the alignment ^17^. By default cognac identifies core genes to a given set of genomes; however, the selection criteria are customizable to allow for flexibility when creating alignments for various applications. The default selection criteria for selecting marker genes are: 1) present in 99% of genomes, 2) present in a single copy in 99.5% of genomes, and 3) ensuring that there are at least two unique alleles for each gene. Allowing some degree of missingness allows for assembly errors which may arise in large datasets.

Once the marker genes are identified, the individual gene alignments of the amino acid sequences for each gene are generated with MAFFT ^18^. To increase efficiency, only unique alleles are aligned. For each gene, duplicated alleles are identified, only the representative unique alleles are input to MAFFT, and the alignment is then parsed to replicate the aligned sequence for each of the duplicated alleles to generate the full alignment for the dataset. Minimizing the number of sequences reduces the computational overhead associated with aligning a large number of sequences, leading to significantly reduced memory consumption and run-time.

Finally, the individual genes are concatenated into a single alignment to be used in downstream analysis. Additionally, several optional outputs may be generated. We provide the functionality to reverse translate the amino acid alignment to return the nucleotide alignment. We use gap placement in the amino acid alignment to position the corresponding codons from the nucleotide sequence of each gene, generating a codon aware nucleotide alignment. This has the added benefit of increasing the number of variant positions in the alignment, which are a product of synonymous substitutions. This is potentially useful for applications where maximizing variation is key. We also provide functionality for parsing the alignments including: eliminating gap positions, removing non-variant positions, partitioning the alignment into the individual gene alignments, removing low quality alignment positions, creating distance matrices, and creating neighbor joining trees within our R package.

### Benchmarking

To demonstrate the utility of our tool, we created genus-level core gene alignments for 27,529 genomes from eight clinically relevant species of bacteria (Table 1, Supplementary table 1). The number of genomes included from each genera had a wide range from 24 for *Pluralibacter* to 11,639 for *Escherichia*. Cognac was run requiring that at least 1000 genes were included in the alignment, iteratively removing genomes that are missing the most frequently observed genes until there are the specified number of core genes included in a set of genomes. Optionally, genomes missing above a threshold of missing genes, 1% by default, are excluded from the analysis. Because this was a large data set with the potential for genomes to be mislabeled, or of poor quality this method ensures that mislabeled genomes do not contribute to the selection of core genes. Additionally, for our test runs we included the optional steps to reverse translate the alignment, create a pairwise single nucleotide variant distance matrix from the nucleotide alignment, and generate a neighbor joining tree.

**Table 1.**
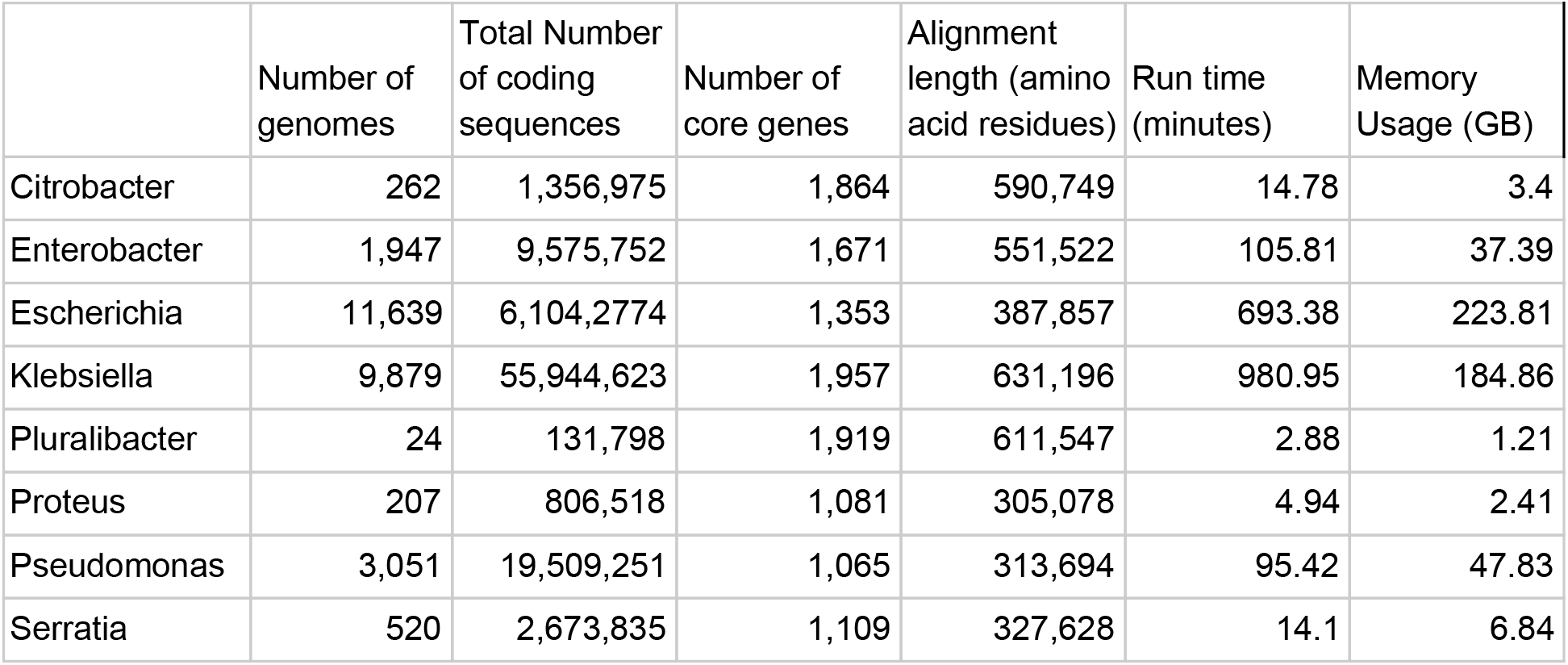
Description of dataset and run statistics for the analysis in this study.

|  | Number of genomes | Total Number of coding sequences | Number of core genes | Alignment length (amino acid residues) | Run time (minutes) | Memory Usage (GB) |
| --- | --- | --- | --- | --- | --- | --- |
| Citrobacter | 262 | 1,356,975 | 1,864 | 590,749 | 14.78 | 3.4 |
| Enterobacter | 1,947 | 9,575,752 | 1,671 | 551,522 | 105.81 | 37.39 |
| Escherichia | 11,639 | 6,104,2774 | 1,353 | 387,857 | 693.38 | 223.81 |
| Klebsiella | 9,879 | 55,944,623 | 1,957 | 631,196 | 980.95 | 184.86 |
| Pluralibacter | 24 | 131,798 | 1,919 | 611,547 | 2.88 | 1.21 |
| Proteus | 207 | 806,518 | 1,081 | 305,078 | 4.94 | 2.41 |
| Pseudomonas | 3,051 | 19,509,251 | 1,065 | 313,694 | 95.42 | 47.83 |
| Serratia | 520 | 2,673,835 | 1,109 | 327,628 | 14.1 | 6.84 |

All runs finished in less than a day, and ranged from three minutes to 16 hours and 21 minutes (Table 1). Run-time grew linearly as the number of genomes increased (Fig. 2a). For all runs, with the exception of Pseudomonas, generating the MAFFT alignments was the largest portion of the total run-time (Fig. 2b). The CD-HIT step was the highest fraction of runtime for Pseudomonas due to the larger genome size and the large degree of pan-genome diversity observed for this genus (Table 1).

**Fig. 2.**
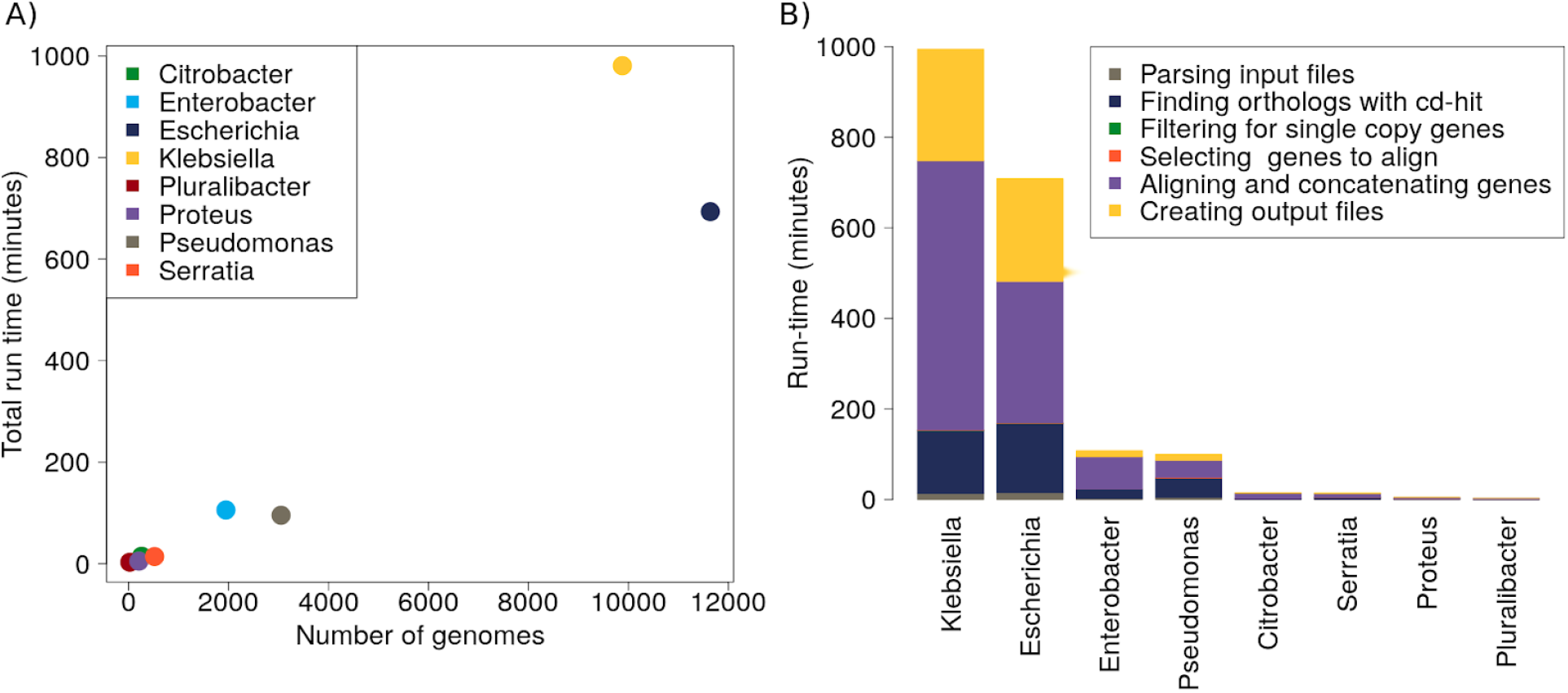
cognac is able to maintain reasonable run time even for very large datasets, generating the alignment, reverse translation, creating a distance matrix, and neighbor joining tree. **a** For each genus the run time plotted against the number of genomes included in the analysis. **b** The composition of the run time by step.

To assess the magnitude of the reduction in the quantity of sequences that were aligned by selecting only the unique alleles of each gene, which is related to increased computational efficiency, we calculated the number of unique alleles per core gene as a fraction of the number of genomes (Table 1, Fig. 3a). We observed a strong inverse relationship between the number of genomes included and the number of unique alleles identified within the dataset (Fig. 3b). As a fraction of the number of genomes, *Klebsiella* had the lowest range of unique alleles with 0.02% (n = 2) to 6.07% (n = 600), with a median of 1.13% (n = 112). Pluralibacter had the fewest genomes and had the highest proportion of unique alleles, with a maximum value of 79.9% unique alleles (n = 19). This is a substantial decrease in the quantity of sequences that need to be aligned, enabling cognac to scale to very large datasets. Because organisms are related genealogically, sequences in the genome are not independent, sharing a common ancestor. Therefore adding additional genomes does not necessarily expand the number of unique alleles for any genes, and all of the sequences may be represented by a substantially reduced subset of the number of samples.

**Fig. 3.**
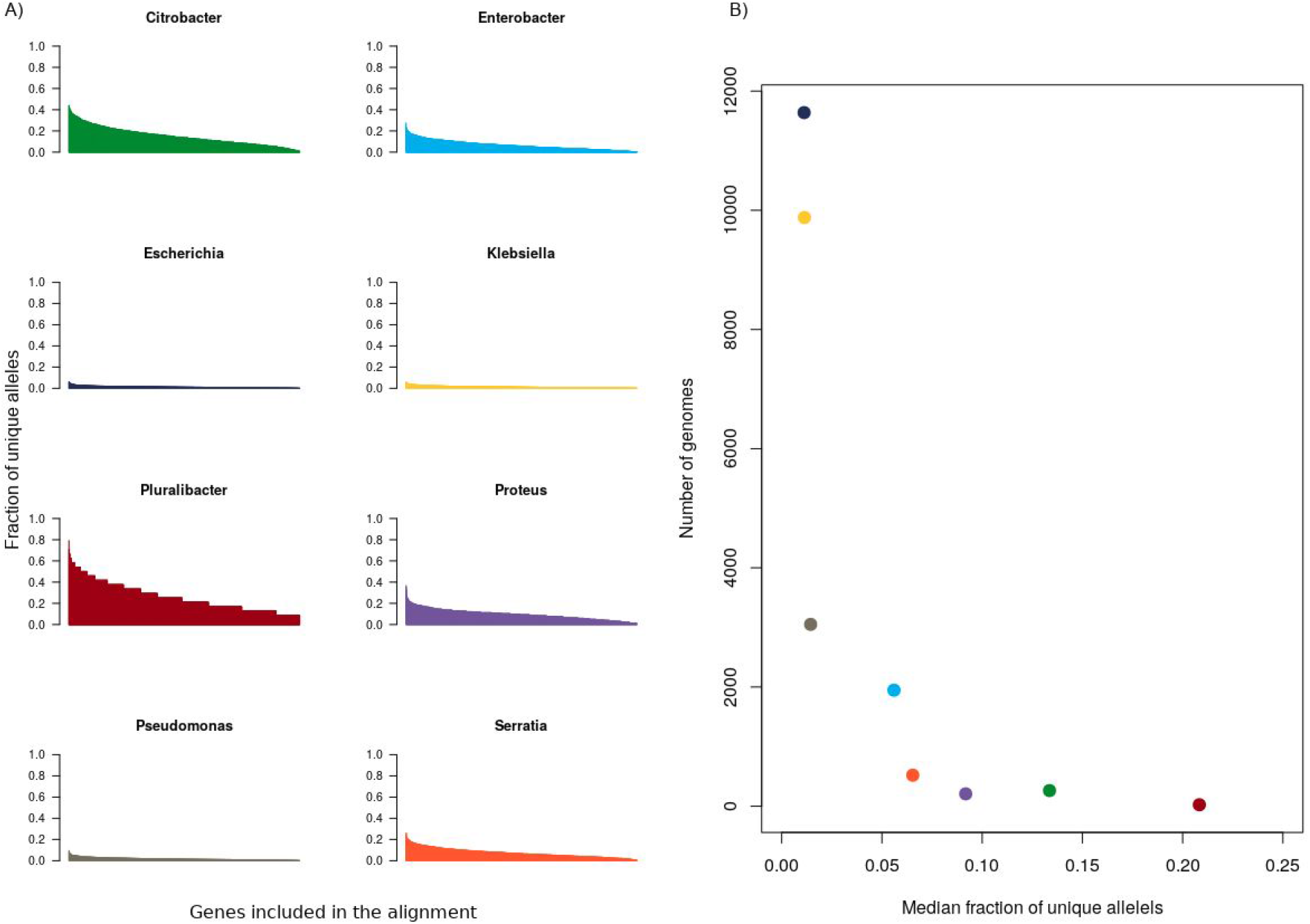
The fraction of unique alleles per gene is inversely proportional to the number of genomes in the dataset. **a** The distribution of unique alleles per core gene included in the alignment as a fraction of the number of genomes. **b** The relationship between the number of genomes and the median fraction of unique alleles for each gene.

We then wanted to analyze the impact of reverse translation in amplifying the sequence diversity in the alignment. The raw number of pairwise substitutions was calculated for each genome from the nucleotide and amino acid alignment (Fig. 4). Reverse translation greatly expanded the quantity of genetic variation contained in the alignment, although to different degrees for different datasets. This may reflect non-biological processes. For example, different data sets may have more diversity due to non-random sampling of the diversity within each genus. Additionally, the magnitude of the phylogenetic distances between isolates may not be uniform within different taxonomic assignments. Although biological factors may also play a role in the observed genetic distances. For example, the lowest amount of diversity was observed in *Pseudomonas*. The published mutation rate for *E. coli* is 2.5 times higher than that of *P. aeruginosa,* suggesting that the differences in diversity may be a function of the mutation rate in these organisms ^19^.

**Fig. 4.**
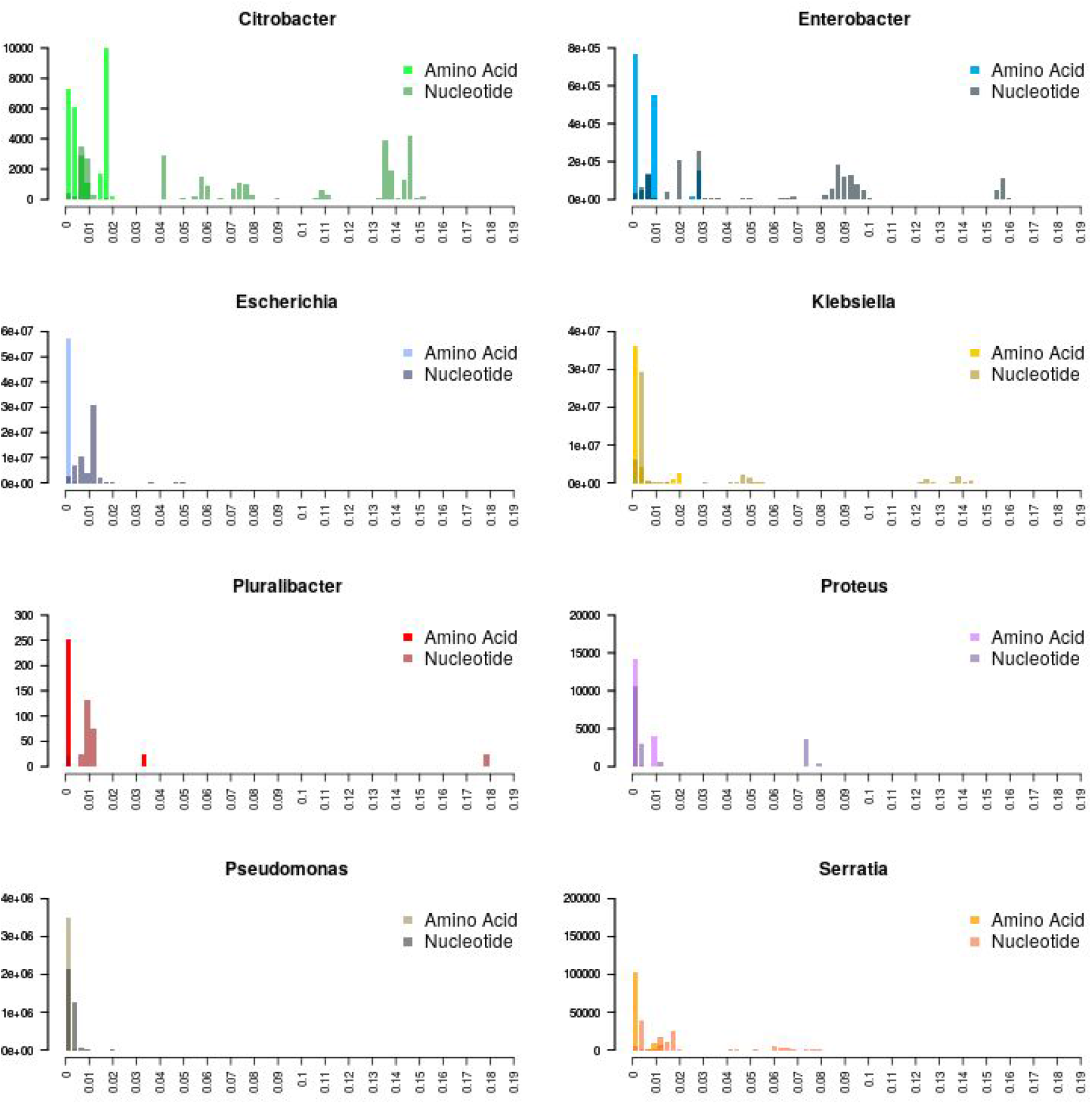
Reverse translation of core gene alignments expands the number of variants for phylogenetic analysis. Pairwise distance matrices were constructed with the raw number of substitutions of the amino acid and nucleotide alignments. Histograms show the distribution of substitutions per position in the alignment. Lighter colors represent the amino acid distances and darker colors represent the nucleotide distances

## Discussion

We present a method to rapidly identify over 1000 marker genes and generate concatenated gene alignments that is capable of handling diverse bacterial genomes. Recently, we used this method to generate a core genome alignment and maximum likelihood tree for 52 genomes in the family *Bacteroidetes*, illustrating the utility of this tool to create gene trees over large phylogenetic distances ^20^. Importantly, phylogenetically informative marker genes are selected using a data driven approach, without any knowledge of the input genomes *a priori*, which allows for flexible selection of marker genes that are tailored to any input dataset.

Our approach relies fundamentally on amino acid sequence comparisons. Translation provides a natural compression algorithm, which has several advantages. First, the amino acid sequences have a third of the length of the corresponding nucleotide sequence. Because the length of the input sequences is a major contributor to the computational complexity of MSA, this reduction in length significantly improves performance and scalability ^21,22^. Additionally, amino acid sequences have a higher degree of conservation relative to nucleotide sequences ^23^. This enables us to leverage redundancy in the codon code to more accurately identify orthologous genes and generate more accurate alignments. This enables a more robust and rapid approach for identifying and aligning orthologous genes, especially when applied to phylogenetically diverse datasets.

When performing computationally intensive procedures, amino acid sequences have many advantages; however, nucleotide alignments may be preferable for some applications. To address this, we provide the optional functionality to reverse translate the amino acid alignment to return the nucleotide alignment. This can substantially increase the sequence variation contained in the alignment, which may be useful for applications where it is important to distinguish between closely related isolates. Additionally, we leverage the information contained in the amino acid sequences to produce a codon aware alignment. This allows for greater accuracy in placement of functional residues within the gene sequence and reduces the potential for misalignment of codons that may occur when aligning nucleotide sequences.

An important feature of our algorithm is that it relies only on whole genome assemblies, which provides several advantages over commonly used techniques of aligning raw sequencing reads to a reference genome. First, with respect to the size of the files, assemblies are a small fraction of the files containing the raw sequencing data. Second, cognac does not require selection of a reference genome. Different choices of reference genome have been shown to have large influences on the quality of the output alignment, potentially amplifying the frequency of mapping errors ^24^. Additionally, the mapping accuracy is severely compromised when considering diverse datasets, even within a species. This limits the application of this method to diverse datasets. Finally, since our approach relies on assemblies, this enables us to analyze genomes sequenced on different platforms, allowing for increased sample size.

Other assembly based methods for estimating the genomic distance between genomes use dimensionality reduction techniques such as k-mers or the MinHash algorithm to estimate the distance between genomes ^25,26^. These methods have the advantage that they can leverage non-coding regions as a source of additional variation; however, the natural structure of the data is lost. Our method not only allows for an estimation of the genetic distances between isolates, but also produces an alignment that can be used in downstream applications. This has the potential to leverage the alignment to identify recombinogenic genes, and has the potential to be used to gain biological insights into molecular evolution.

Our algorithm was able to scale to extremely large datasets. For a data set of 11,639 Escherichia genomes we were able to generate a neighbor joining tree from a nucleotide concatenated gene alignment in less than 17 hours. This is accomplished by reducing the computational overhead of MSA in two ways: 1) translating the sequences, effectively reducing their length; and 2) reducing the number of sequences by only aligning unique alleles. For extremely large datasets, this results in an approximately 99% reduction in the number of sequences that need to be aligned, allowing for great improvements in scalability, and allows for application to extremely large datasets.

## Conclusions

In summary, cognac is a robust, rapid method for generating concatenated gene alignments that scales to extremely large datasets. Our method uses a data driven approach for identification of phylogenetic markers, which are efficiently aligned and concatenated into a single alignment for downstream phylogenetic analysis. The pipeline is open source and freely available as an R package. We expect our tool will be generally useful for many different types of analysis and will enable evolutionary insights in a broad range of applications.

## Methods

Genomes for this study were collected as part of a longitudinal study of carbapenem resistant organisms and additional genomes were downloaded from the Pathosystems Resource Integration Center (PATRIC) ^27,28^. All available genomes from PATRIC as of 06/01/2020 that were isolated from humans and met the criteria for good quality were downloaded from the PATRIC FTP server. Quality was assessed for completeness, contamination, coarse consistency, and fine consistency via the CheckM algorithm within the PATRIC genome annotation service ^29,30^.

Cognac was developed for R version 3.6.1. C++ was integrated via the Rcpp package (version 1.0.3) and was written using the C++11 standard ^31^. Multithreading is enabled in the C++ code via RcppParallel, which provides wrapper classes for R objects used by Intel Threading Building Blocks parallel computing library ^32^. Multithreading for R functions was enabled via the future.apply package (version 1.3.0) ^33^. Functions for analysis of phylogenetic trees were enabled via the APE R package (version 5.3).^34^ Clustering genes is performed with CD-HIT (version 4.7) and alignments are generated via MAFFT (v7.310) ^17,18^.

## Availability of data and materials

cognac source code is available at https://github.com/rdcrawford/cognac under a GNU General Public License, version 2. Whole genome sequences and annotations used in this study are available from https://www.patricbrc.org/ and RefSeq under BioProject PRJNA603790 and PRJNAXXXXXX. Scripts used in benchmarking are available at https://github.com/rdcrawford/cognac_paper.

